# Unraveling yeast diversity in food fermentation using ITS1-2 amplicon-based metabarcoding

**DOI:** 10.1101/2025.11.05.686804

**Authors:** Ines Pradal, Thomas Gettemans, Stefan Weckx

## Abstract

A detailed characterization of the microbial ecosystem involved in the production processes of fermented foods is essential. Although fermented foods are an important part of the human diet and have seen an increasing interest nowadays, some challenges still need to be solved. Specifically, yeast identification through culture-independent methodologies is still limited to the genus level. Unlike for bacterial species identifications, long-read sequencing technologies have barely been used for yeast species identification, and, to the best of the authors’ knowledge, it has not been validated with mock communities reflecting food fermentation processes yet. Therefore, in the current study, we present an amplicon-based metabarcoding approach targeting the full-length internal transcribed spacer (ITS) region comprising the ITS1, 5.8S rRNA gene, and ITS2 using the PacBio HiFi sequencing platform. This method was validated using mock communities composed of yeast species involved in sourdough, lambic beer, and cocoa fermentation processes. Accurate species-level identification was achieved for most of the species. However, special attention should be given to *Saccharomyces*-rich niches, as accurate species-level identification for this genus is still challenging. Furthermore, underestimation of the relative abundance of species with short ITS regions, such as *Pichia* and *Brettanomyces*, occurred. In addition, the method was successfully applied to describe the yeast diversity present in two sourdough and two lambic beer samples. Overall, the current method provides an unprecedented way of determining the species-level yeast composition of complex ecosystems present in fermented food products.

**Importance:** To date, species-level identification of common yeasts present in food fermentation ecosystems has been difficult, if not impossible, when using short-read sequencing methods. However, species-level identification is essential when evaluating and describing the characteristics of fermented food microbiomes. The current study reports on the development and validation of an amplicon-based metabarcoding approach combined with long-read PacBio HiFi sequencing targeting the full ITS region, comprising the ITS1 and ITS2 regions as well as the 5.8S rRNA gene. The described methodology enables species-level identification of the most common yeasts present in food fermentation ecosystems. This new methodology is of importance for all researchers in the field of fermented foods. By extension, researchers in other fields of microbiology can find inspiration in this paper.

## 1. Introduction

Fermented foods and beverages have been an important part of the human diet for centuries and have seen an increasing interest nowadays (1). As a result of the great importance of fermented foods and beverages, studying the microbial ecosystem composition involved in their production processes in detail is essential. Fermented foods are produced through the metabolic activity of yeasts, lactic acid bacteria (LAB), coagulase-negative cocci, acetic acid bacteria (AAB), and/or filamentous fungi (1–3). Among the microorganisms involved, yeasts are key microorganisms during the production of sourdough, beer, and wine, among others, as well as during cocoa fermentation (2, 4).

Yeast diversity of food fermentation processes has been traditionally studied by culture-dependent approaches in the same way as is done for the bacterial diversity (5). However, besides the benefit of being able to construct a collection of microbial strains for further investigation, those approaches are time-consuming and are biased by the selective growth media and conditions used (6, 5). The advances in high-throughput sequencing technologies allow the study of microbial communities by using a culture-independent approach (7, 5). Specifically, amplicon-based metabarcoding, also known as metagenetics, has been the gold standard in this field, as it has a high throughput with relatively low costs (7–11). This technique consists of the amplification of a specific genomic region expected to be present in all genomes of the microorganisms present in the ecosystem, followed by the sequencing of the amplicons obtained (7). Therefore, selecting an appropriate genomic region that enables a good taxonomic resolution is of crucial importance. Furthermore, selecting a proper sequencing technology is also of importance, as the technology chosen can also impose certain constraints. For example, Illumina MiSeq and Illumina NovaSeq are the most commonly used technologies, but they only allow for sequencing of short reads, despite their high accuracy. This short length results in a taxonomical resolution that seldom reaches the species level (10, 12). Nevertheless, nowadays, long-read sequencing technologies, such as Pacific Biosciences’ (PacBio) HiFi long-read sequencing or Oxford Nanopore Technologies’ nanopore sequencing, are available on the market. The former one has emerged as a suitable technology for amplicon-based metabarcoding as it combines long reads with high accuracy (10, 13).

In the case of bacterial communities, the 16S rRNA gene is commonly considered to be the most suitable gene for identification (14, 15). Due to the length restriction imposed for Illumina sequencing, specific variable regions or combinations thereof, such as V1-V3, V4, and V3-V5, have been targeted (15–17). However, this only allowed for genus-level resolution (17, 18). Recently, the full 16S rRNA gene has been targeted thanks to PacBio’s circular consensus sequencing, nowadays branded as HiFi sequencing (10, 15, 19, 20). This new method allows a species-level resolution for most of the LAB commonly found during food fermentation. Specifically, it has been used to reveal the bacterial diversity of beer (21, 22), cheese (23, 24), and sourdough (25).

In the case of yeasts, there is no such gold standard. Variable regions of the 18S rRNA gene, the internal transcribed spacer 1 (ITS1) region, and the ITS2 region are some examples of the genomic regions targeted (9, 26–28). In all cases, the short length imposed by Illumina sequencing also leads to less accurate genus-level identification compared to long-read amplicon sequencing (29). In addition, comparisons of taxonomic assignment using different regions are available for specific samples, such as beer (27), wine (27), soil (9), human saliva (9), grape must (9), and bioaerosols (30). However, different microbial compositions are retrieved for different targeted regions, and the preferred region is environment-dependent. Thus, no consensus regarding which genomic region to target has been reached yet. Nevertheless, all studies pointed out the ITS regions as the primary fungal barcode because of a higher number of sequence variations (12, 31). Recently, PacBio HiFi sequencing of the genomic region comprising the ITS1, 5.8S rRNA gene, and ITS2 has been performed for metabarcoding of eukaryote-containing samples. Specifically, it has been used to describe the fungal diversity of coffee plant samples (32), wheat, maize, barley and cover crops (clover, hairy vetch, oilseed radish and fallow) samples (33), sea water samples (34), and soil samples (31, 35), with most studies focusing on filamentous fungi rather than yeasts. Nevertheless, this method has been barely reported for food fermentation samples (21), and, to the best of the authors’ knowledge, it has not been validated with mock communities reflecting food fermentation processes yet.

Hence, this study aimed to assess whether amplicon-based metabarcoding targeting the full ITS region, comprising the ITS1 and ITS2 regions as well as the 5.8S rRNA gene, enables species-level identification of the most common yeasts present in food fermentation ecosystems. After determining the appropriate primer sets and amplification conditions, the technique was tested using mock communities representing the yeast composition of different fermentation stages of relevant fermented foods and beverages, namely sourdough, lambic beer, and cocoa. Finally, the fungal diversity of two sourdough samples and two lambic beer samples was assessed using this technique for validation.

## 2. Results

### 2.1. Primer selection and optimization of the PCR conditions: BITS-ITS4 vs ITS1-ITS4

PCR amplification using the BITS-ITS4 primer set (the BITS primer was extended with barcodes 04, 05, and 06; the ITS4 primer was extended with barcodes 19 and 20) resulted in no visible fragments on gel electrophoresis (data not shown). Hence, the primer combination BITS-ITS4 was deemed to be unsuccessful at properly amplifying the targeted microorganisms. On the contrary, the use of the ITS1-ITS4 primer set (Supplementary Table S2) resulted in visually successful amplification. Specifically, the clearest bands were obtained when an annealing temperature of 52 °C was used. The purification of the amplicons obtained resulted in DNA concentrations between 4.9 ng/µl and 49.0 ng/µl. Notably, PCR amplifications involving the ITS1 primer extended with barcode 05 resulted in no amplification, most likely due to secondary structures formed between the primer and barcode sequences. Indeed, when checking the extended primer in OligoEvaluator (https://www.oligoevaluator.com/LoginServlet), four possible moderate secondary structures were present involving the 3’ end of the primer, possibly preventing primer annealing.

### 2.2. Sequencing data

An average of 118,998 reads per sample was obtained with a minimum of 9,718 reads and a maximum of 348,099 reads (Supplementary Table S2). Those reads had an average length of 763 nucleotides, with 90 % of the reads being between 600 and 900 nucleotides, while 10 % of the reads was shorter than 600 nucleotides. Primer removal, quality filtering, and denoising resulted in the removal of 11 % of the reads, yielding an average of 105,470 reads per sample, with a minimum of 8,007 reads (Lambic beer A) and a maximum of 310,563 reads [Mock community (MC) lambic beer equal 1]. Most of the removed reads (8 % of the total reads) were eliminated during the quality filtering step. After data processing, the average length of the reads was 722 nucleotides, with a minimum length of 405 and a maximum length of 842 nucleotides. Only 15 % of the reads were shorter than 600 bp.

Rarefaction analysis showed that all MCs and food fermentation samples were sequenced with a sufficient depth to reliably assess the full fungal diversity (Supplementary Fig. S1). Specifically, in the case of the Lambic beer A and B samples, the plateau was reached with 4,741 and 5,301 reads, respectively, representing a maximum of 17 and six species, respectively. In the case of the Sourdough A and B samples, the plateau was reached with 43,521 and 8,041 reads, respectively, representing a maximum of 30 and 33 species, respectively.

### 2.3. Taxonomic classification

#### 2.3.1. Above genus reads

Taxonomical assignment at the species or genus level could not be achieved for all amplicon sequence variants (ASVs) obtained. The proportion of above genus reads (AGR) was small for the different MCs, with a maximal percentage of 0.2 % of the total reads in the MC cocoa equal 3. The Lambic beer A and Lambic beer B samples displayed similarly low percentages of AGR. On the contrary, the sourdough samples had large amounts of AGR with 16.7 % and 60.8 % in the Sourdough A and Sourdough B, respectively. Blastn searches of these ASVs revealed that 99.4 % and 99.8 % of the AGR, respectively, aligned to the ITS region of *Simplicia felix* (87 - 91 % sequence identity and 85.6 - 88.3 % coverage). Further, blastn searches of these ASVs against the genomes of type material of *Triticum aestivum* (soft wheat) and *Secale cereale* (rye) resulted in alignments of the ASVs with the ITS regions of these two plant species with 100 % sequence identity and coverage. Therefore, the AGR were removed before calculating the relative abundances of fungal species in each sample.

#### 2.3.2. Mock communities

A good reproducibility was found among the three replicates of all MCs (Fig. 1).

**Fig. 1.**
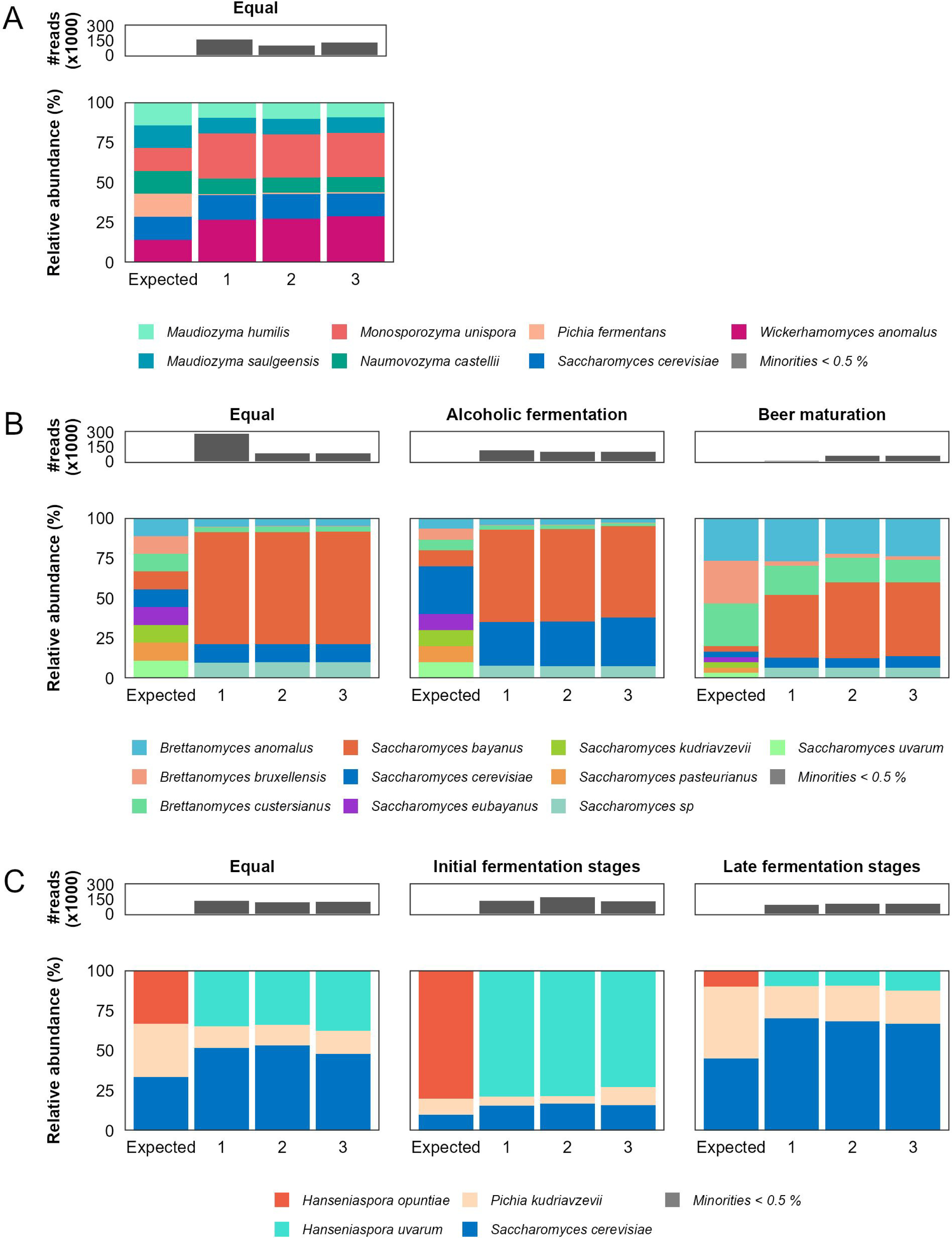
Microbial composition of the sourdough (A), lambic beer (B), and cocoa (C) mock communities, depicting both the expected and obtained relative abundances. Sample numbers 1, 2, and 3 represent the different replicates.

##### Sourdough MC

All species included in the MC were retrieved, corresponding with a precision of 1 (Fig. 1A and Table 2). Nevertheless, the divergence between the expected composition and the obtained one was 0.27, as some species were found in relative abundances different from the expected ones. Specifically, the obtained relative abundance of *Saccharomyces cerevisiae* was very similar to the expected one [on average 0.6 ± 0.7 %points (%pt.) more]. In the case of *Maudiozyma humilis*, *Maudiozyma saulgeensis*, and *Naumovozyma castelli*, their relative abundances were slightly below the expected ones (on average 4.6 ± 0.2 %pt. less). In contrast, *Monosporozyma unispora* and *Wickerhamomyces anomalus* were found at a higher relative abundance than the expected ones (on average 13.3 ± 0.8 %pt. more). Finally, the relative abundance of *Pichia fermentans* was on average 0.8 ± 0.1 %, almost 20 times lower than the expected one.

**Table 1.**
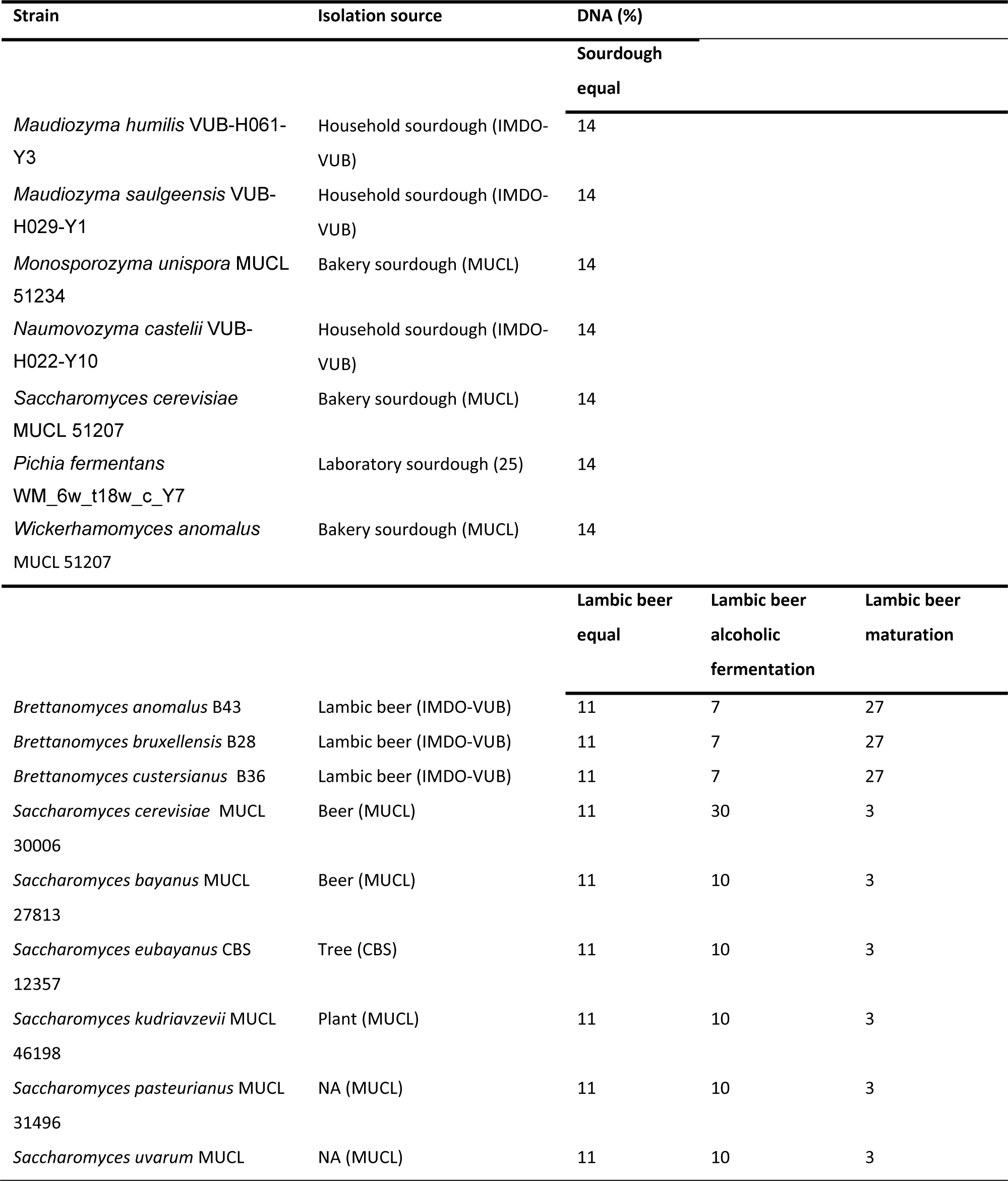
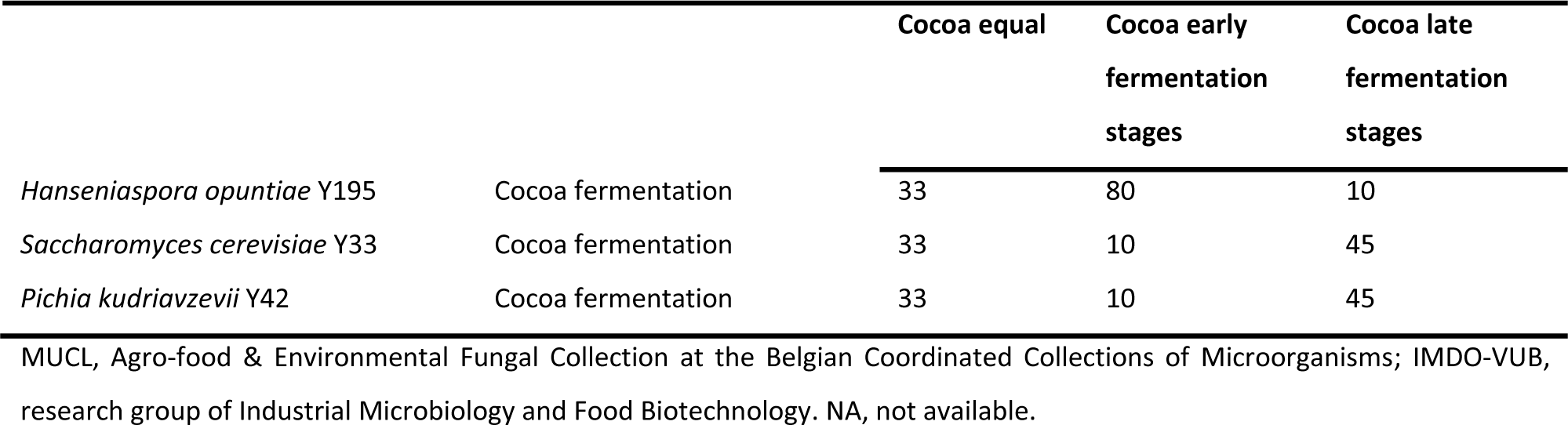
Yeast strains and composition of the mock communities used in the study.

**Table 2.** Metrics describing the obtained deviations from the expected composition of the mock communities.

| Mock community | Divergence | False<br>negatives | False positives | True positives | Precision |
| --- | --- | --- | --- | --- | --- |
| Sourdough equal | 0.2743 ± 0.0051 | 0 | 0 | 7 | 1 |
| Lambic beer equal | 0.6915 ± 0.0006 | 4 | 1 | 5 | 0.83 |
| Lambic beer alcoholic fermentation | 0.5527 ± 0.0016 | 4 | 1 | 5 | 0.83 |
| Lambic beer maturation | 0.5073 ± 0.3595 | 4 | 1 | 5 | 0.83 |
| Cocoa equal | 0.5321 ± 0.0063 | 1 | 1 | 2 | 0.67 |
| Cocoa early fermentation stages | 0.8318 ± 0.0228 | 1 | 1 | 2 | 0.67 |
| Cocoa late fermentation stages | 0.3391 ± 0.0081 | 1 | 1 | 2 | 0.67 |

##### Lambic beer MCs

The composition of the lambic beer MCs was determined with a precision of 0.83 in all cases, and a divergence of, on average, 0.69, 0.55, and 0.51 in the case of the lambic beer equal, lambic beer alcoholic fermentation, and lambic beer maturation MCs, respectively (Table 2). A precision lower than 1 could be explained by the fact that the species *Saccharomyces eubayanus*, *Saccharomyces kudriavzevii*, *Saccharomyces pasteurianus*, and *Saccharomyces uvarum* were not found in any of the lambic beer MCs (Fig. 1B). Remarkably, 10.0 ± 0.0, 7.7 ± 0.1, and 6.3 ± 0.2 % of the reads of the lambic beer equal, lambic beer alcoholic fermentation, and lambic beer maturation MCs, respectively, were assigned to *Saccharomyces* sp. Blastn searches using the ASVs assigned to this taxonomic unit as a query resulted in the alignment of seven out of eight ASVs to *S. kudriavzevii* with at least 98.4 % sequence identity and 100 % coverage. In contrast, the eighth ASV aligned to *S. kudriavzevii* and *Saccharomyces paradoxus* with a sequence identity of 95.8 % and a coverage of 100 %. As was the case for the sourdough MC, *S*. *cerevisiae* was found at a similar relative abundance to the expected ones in the lambic beer equal (on average 11.4 ± 0.1 *vs.* 11.1 %) and the lambic beer alcoholic fermentation MCs (on average 28.7 ± 1.4 *vs.* 30.0 %). However, this species was found at a relative abundance of 6.5 ± 0.4 %, compared to the expected 3.3 % in the lambic beer maturation MCs. *Saccharomyces bayanus* was found at a higher relative abundance than those expected in the lambic beer equal (70.1 ± 0.1 *vs.* 11.1 %), lambic beer alcoholic fermentation (57.4 ± 0.2 *vs.* 10.0 %), and lambic beer maturation (44.2 ± 3.6 *vs.* 3.3 %) MCs. Furthermore, the *Brettanomyces* species were all underrepresented in the three lambic beer MCs, with the case of *Brettanomyces bruxellensis* being the most remarkable one (relative abundances lower than 2.4 % in all cases).

##### Cocoa MCs

The cocoa MCs were described with a precision of 0.67, and divergences of 0.53, 0.83, and 0.34 in the case of the cocoa equal, cocoa early fermentation stages, and cocoa late fermentation stages MCs, respectively (Table 2). These values were the result of incorrect identification of the *Hanseniaspora* species, leading to higher divergences for the MCs with higher expected relative abundances for *Hanseniaspora*. Specifically, reads from *Hanseniaspora opuntiae* were identified as *Hanseniaspora uvarum*. Nevertheless, its relative abundances were very similar to the expected ones in the cocoa equal (35.5 ± 1.7 *vs.* 33.3 %), cocoa early fermentation stages (75.5 ± 2.8 *vs.* 80 %), and cocoa late fermentation stages (10.6 ± 1.4 *vs.* 10 %) MCs. *S. cerevisiae* was found with a relative abundance 1.5 times higher than the expected one in all cocoa MCs (Fig. 1C). In contrast, *Pichia kudriavzevii* was found with a relative abundance that was 2.5, 1.4, and 2.1 times lower than the expected ones for the equal cocoa, cocoa early fermentation stages, and cocoa late fermentation stages MCs, respectively.

#### 2.3.3. Food fermentation samples

When applying the developed method to the food fermentation samples, a high diversity of yeast species was retrieved (Fig. 2). The lambic beer samples, Lambic beer A and Lambic beer B, were both mostly inhabited by *S. cerevisiae* (Relative abundance of 76.8 % and 50.8 %, respectively), *S. bayanus* (10.3 % and 19.8 %, respectively), and *Brettanomyces anomalus* (3.0 % and 29.1 %, respectively). The sourdough samples displayed different yeast compositions with Sourdough A containing *Maudiozyma pseudohumilis* (90.1 %), *Debaryomyces prosopidis* (3.0 %) and *S. cerevisiae* (1.9 %) and Sourdough B containing *Ma. humilis* (47.9 %), *D. prosopidis* (27.7 %) and *S. cerevisiae* (11.2 %). In the sourdough samples, two filamentous fungi were retrieved at low relative abundances, namely, *Alternaria* sp. and *Cladosporium herbarum*.

**Fig. 2.**
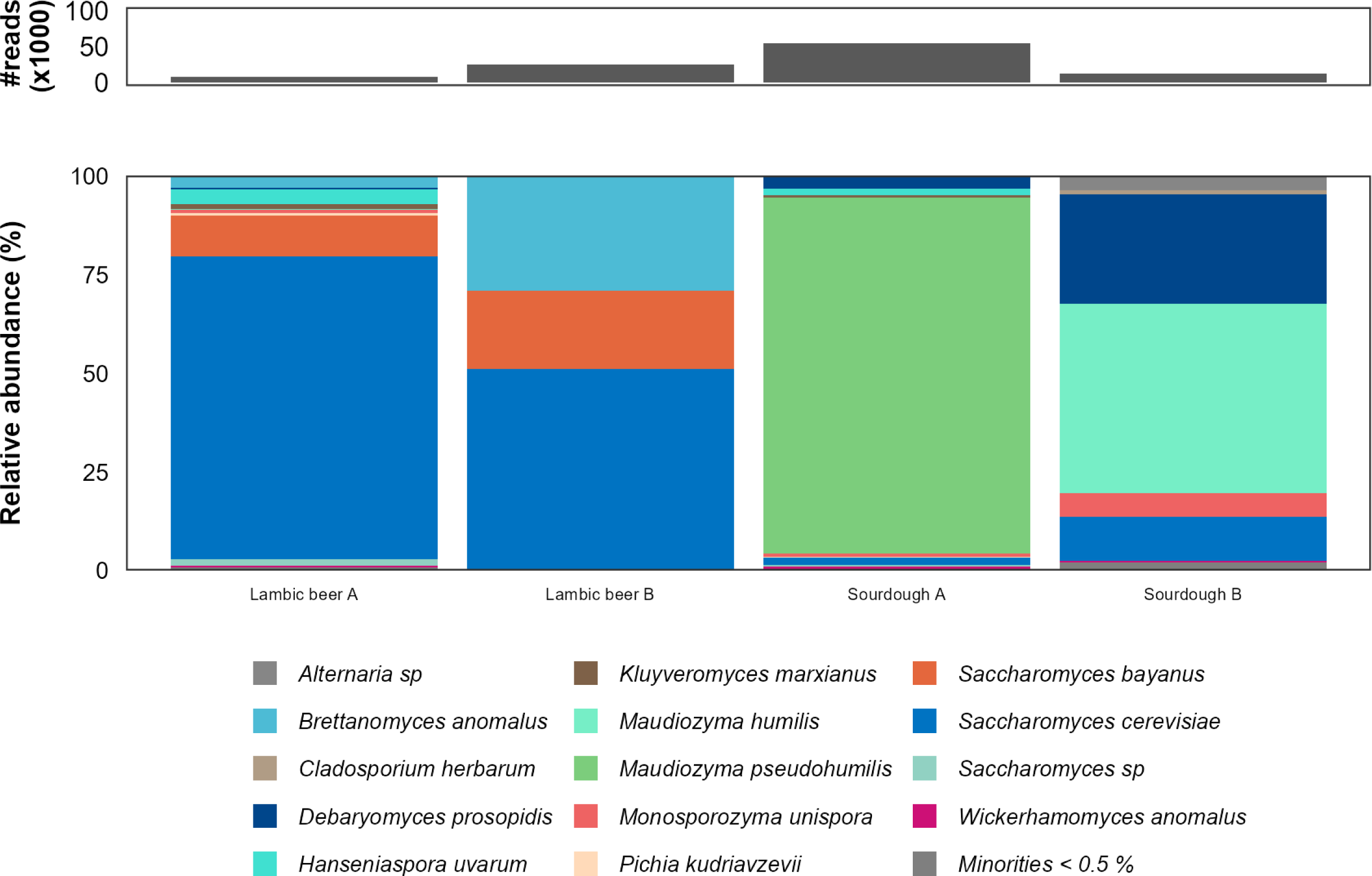
Microbial composition of the food fermentation samples.

## 3. Discussion

In the present study, the potential of an amplicon-based metabarcoding method targeting the full ITS region to reveal the yeast diversity of food fermentation samples was investigated. Up to now, culture-independent approaches revealing the yeast diversity in food fermentations have been performed using short-read sequencing, targeting a small region of the fungal rRNA operon, resulting in an identification that was limited at the genus level (22, 25, 27, 36–38). However, the contribution to food fermentation processes of different species within a genus might differ. Therefore, targeting a larger region that would allow species-level identification was needed in the field.

Thanks to PacBio’s long-read HiFi sequencing technology, in combination with PacBio’s Kinnex library construction strategy, longer amplicons can be sequenced in a cost-efficient manner, providing a higher level of taxonomic resolution compared to previous methods. Whereas the PacBio-based method has already been used to sequence the full ITS region to investigate fungal diversity, those studies focused in most cases on filamentous fungi, and not on yeasts (21, 31, 33–35). Whether this method would be applicable in the field of food fermentation, in which yeasts are key microorganisms (1–3), remained unclear.

As a first step, seven DNA-based mock communities were composed, representing different stages of sourdough, lambic beer, and cocoa fermentation processes. These mock communities were composed of different species of the same genus, including *B. anomalus, B. bruxellensis, Brettanomyces custersianus*, *Ma. humilis*, *Ma. saulgeensis*, *Mo. unispora*, *N. castelli, S. cerevisiae*, *P. fermentans, P. kudriavzevii*, and *W. anomalus*. Overall, species-level identification was successfully obtained. However, the largest discrepancy was the misidentification of *H. opuntiae* as *H. uvarum*, which is a known issue and is the result of their close relationship and highly similar ITS regions, despite them being different species (39). Furthermore, the taxonomic resolution at the species level within the *Saccharomyces* genus using the ITS region is known to be limited, as several species other than *S. cerevisiae* have identical sequences of this region (40). Therefore, special attention should be given to the interpretation of the results in the case that *Saccharomyces*-rich niches are investigated.

The lack of species-level resolution or species misidentifications, as for the specific taxa discussed above, could be solved by targeting longer regions, such as the full rRNA operon, spanning the 18S rRNA gene, ITS1, the 5.8S rRNA gene, ITS2, and the 28S rRNA gene (41, 42). However, such an approach would imply other challenges. Although the length of the expected amplicon, 6 kilobases (kb), fits perfectly within PacBio’s HiFi and Kinnex sequencing strategies, well-curated, publicly available databases containing the full operon are currently not available. This could be overcome by the construction of custom databases (41, 42). However, the need to construct such custom databases as well as perform regular updates could prevent this strategy from becoming a routine strategy in microbiology laboratories unspecialized in advanced bioinformatics. Nevertheless, the launch of the Ribosomal Operon Database (43) might change this scenario, although a regular update of such a recent database cannot be taken for granted, which, given the regular reclassifications of yeasts in recent years, is crucial for accurate yeast identification (44–48).

In addition to accurate species-level identification, the quantitative estimation of the fungal composition, expressed as relative abundance, should be as accurate as possible to obtain a good representation of the real community. However, this is not always possible, as different biases play a role that are difficult (if not impossible) to avoid. In particular, the lower-than-expected relative abundances of *Pichia* species in the sourdough and cocoa MCs are in line with previous studies (22, 25). Due to the short ITS regions for both *Pichia* and *Brettanomyces* (approx. 400-500 bp compared to 700-800 bp for *Saccharomyces*), the underestimation of *Brettanomyces* species in the lambic beer MCs might have been caused by the same effects. However, further investigation is needed, as length biases do not seem to have a great impact on the relative abundances when targeting either the ITS1 or ITS2 region (12). In addition, the different copy numbers of the ITS region in different yeasts might have introduced a bias during the PCR amplification. However, information about exact yeast ITS region copy numbers is still scarce and seems to be strain-dependent (43).

Given the promising results of the MCs, the use of the newly developed method on sourdough and lambic beer samples resulted in a confident description of the fungal diversity at the species level using a culture-independent approach. The species corresponded with those generally described for these matrices using a culture-dependent approach (49, 50). Nevertheless, special attention should be paid to *S. bayanus* for lambic beer samples, as other *Saccharomyces* species might also be present. Furthermore, the high level of AGR in the sourdough samples could be explained by the presence of the ITS region in plants, which in some cases might be similar enough to allow primer hybridization, as described before (12, 51). Therefore, when using this method in cereal-rich matrices, a high sequencing depth should be ensured to have a sufficient number of sequence reads to be able to reliably describe the full yeast diversity after the removal of the AGR, as was the case in the present study.

In conclusion, amplicon-based metabarcoding targeting the full ITS region using the PacBio HiFi sequencing technology is a promising method to unravel yeast diversity in food fermentation samples, achieving a better taxonomic resolution compared to the commonly used Illumina-based methodologies. However, further research is needed to find ways to improve the identification at the species level in environments rich in *Saccharomyces* and *Hanseniaspora* species.

## 4. Materials and methods

### 4.1. Mock communities

To critically assess the methodology developed during the present study that aimed to investigate yeast diversity through amplicon-based metabarcoding of the ITS1-2 region, different DNA-based MCs were constructed representing strains of species that play an important role in food fermentation processes.

#### 4.1.1. Strains, growth conditions, and yeast genomic DNA extraction

All yeast strains used in the present study (Table 1) were stored at -80 °C in cryovials containing yeast extract-peptone-glucose (YPG; Oxoid, Basingstoke, Hampshire, United Kingdom) cultures (52), supplemented with 25 % (v/v) glycerol (Sigma-Aldrich, Saint-Louis, Missouri, USA) as cryoprotectant, as part of the laboratory collection of the research group of Industrial Microbiology and Food Biotechnology (IMDO). Each yeast strain was streaked on YPG agar medium supplemented with 200 ppm of chloramphenicol (Sigma-Aldrich) and incubated at 30 °C for 48 h. A single colony of each strain was transferred to 10 ml of liquid YPG medium and incubated at the same temperature for 24 h. Yeast cell pellets were obtained by centrifugation of 2 ml of the latter cultures at 21,100 × *g* for 5 min. Yeast genomic DNA was extracted from these cell pellets using 600 µl of enzymatic lysis buffer with lyticase (200 U; Sigma-Aldrich), zymolyase (15 U; G-Biosciences, Saint Louis, Missouri, USA), and proteinase K (60 mAnsonU; Merck, Darmstadt, Germany) as described before (24). DNA purification was performed using a DNeasy Blood and Tissue kit (Qiagen, Hilden, Germany). The DNA concentrations were measured by fluorimetry (Qubit; Thermo Fisher Scientific, Waltham, MA, USA).

#### 4.2.2. Composition of the mock communities

A total of seven different MC representing common yeast species present in the microbial communities of various food fermentation processes, namely those of sourdough, lambic beer, and cocoa, were constructed (Table 1). In the case of sourdough, one MC containing equal concentrations of DNA of seven common sourdough yeast species (50, 53) was constructed. In the case of lambic beer, three MCs were considered, namely one MC with equal concentrations of DNA of three *Brettanomyces* and six *Saccharomyces* species, one MC representing the alcoholic fermentation phase of lambic beer that is rich in *Saccharomyces* species, and one MC representing the maturation phase of lambic beer that is rich in *Brettanomyces* species (49). In the case of cocoa, three MCs were considered, namely, one MC with equal concentrations of DNA of *H. opuntiae*, *S. cerevisiae* and *P. kudriavzevii*, one MC representing the early stage of the cocoa fermentation process that is rich in *Hanseniaspora* species (4), and one MC representing the late stage of the cocoa fermentation process that is rich in *S. cerevisiae* and *Pichia* species (4). Each MC was constructed by pooling the strains’ DNA (Table 1) to get a final total DNA concentration of 1.5 ng/µl.

### 4.3. Food fermentation samples

#### 4.3.1. Sampling

To assess the method’s validity in food fermentation samples, two sourdough samples and two lambic beer samples were analyzed. The sourdough samples were both spontaneous Type 1 sourdoughs originating from two artisan bakeries, with Sourdough A being a three-year-old rye sourdough and Sourdough B being a nine-year-old wheat sourdough. Cell pellets were collected by diluting the sourdoughs 1:10 with sterile peptone saline [0.1 % bacteriological peptone (Oxoid), 0.85 % NaCl (Merck)] as described before (25). After mixing to get a homogenous sample, a two-step centrifugation was performed. First, the homogenous samples were centrifuged at 1000 x *g* for 5 min. Second, the flour debris-free supernatants were centrifuged at 5000 x *g* for 20 min to collect the cell pellet. The lambic beer samples consisted of samples taken during the production of lambic beer, with Lambic beer A taken after five days of barrel fermentation and Lambic beer B taken after three months of barrel fermentation. The cell pellets were collected by a centrifugation step at 5000 x *g* for 20 min (21). In both cases, the cell pellets were stored at -20 °C until further DNA extraction.

#### 4.3.2. DNA extraction from food fermentation samples

Extraction of total DNA from the cell pellets obtained was performed as described before for sourdough (25) and lambic beer samples (22). Briefly, in both cases, a combination of both a bacterial lysis solution [300 µl of bacterial lysis buffer with 100 U mutanolysin (Sigma-Aldrich) and 320 kU of lysozyme (Merck)] and a yeast lysis solution [600 µl of yeast lysis buffer with 15 U zymolyase (G-biosciences) and 200 U lyticase (Sigma-Aldrich)] was used to lyse the bacterial and yeast cells. Further, mechanical lysis was performed using UV-C-irradiated glass beads (Sigma-Aldrich). Then, protein digestion was performed with 40.0 μl of a 20.0 % (m/m) sodium dodecyl sulphate (SDS; Sigma-Aldrich) solution and 50.0 μl of proteinase K solution (2.0 mg/ml of proteinase K; Merck) per sample. In the case of the sourdough samples, they were further incubated in a 5.0 M NaCl (Merck) solution and a 10.0-% (m/m) cetyltrimethylammonium bromide (CTAB; Merck) solution. Finally, DNA was purified using chloroform-phenol-isoamyl alcohol solution and a DNeasy Blood and Tissue kit (Qiagen). The DNA concentrations were measured by Qubit fluorimetry (Thermo Fisher Scientific).

### 4.4. PCR

#### 4.4.1. Selection of the primer sets

From all primers described to target the yeast rRNA operon (27, 54, 55), primers BITS (5’-ACCTGCGGARGGATCA-3’) and ITS1 (5’-TCCGTAGGTGAACCTGCGG-3’) were selected as forward primers, and primer ITS4 (5’-TCCTCCGCTTATTGATATGC-3’) was selected as reverse primer. The two resulting primer pairs were selected as they allow the amplification of the full internal transcribed region (ITS1-2). Specifically, the BITS primer - together with B58S3 as a reverse primer - has been used to target the fungal ITS1 region to describe the fungal diversity of food fermentation processes using Illumina amplicon-based metabarcoding (22, 25, 27, 36–38). The primers ITS1 and ITS4 are generally used to amplify the ITS1-2 region to identify yeast isolates in a culture-dependent approach (22, 25, 37, 38, 56, 57). The primers were tagged with Kinnex adaptors and 5’ sample-specific barcodes (Integrated DNA Technologies, Leuven, Belgium; Supplementary Table S1) to allow for multiplexed sequencing, according to the manufacturer’s instructions (Pacific Biosciences, Menlo Park, CA, United States).

#### 4.4.2. Temperature gradient

PCR assays were performed using KAPA HiFi DNA Polymerase (Hot Start and Ready Mix; Roche, Basel, Switzerland) based on the PCR assay described for 16S rRNA gene amplification for PacBio HiFi sequencing (24), but with a different annealing temperature. Specifically, per sample, 1.5 µL of DNase and RNase-free water (VWR International, Radnor, PA, USA), 12.5 µL of 2X KAPA HiFi HotStart ReadyMix (Roche), 3 µl of the forward primer and 3 µl of the reverse primers (2.5 µM each), and 5 µl of template DNA (1.5 ng/µl) were mixed. To assess the most optimal annealing temperature, PCR assays were performed applying a temperature gradient using a TProfessional Basic Gradient thermocycler (BioMetra, Gottingen, Germany) with values 52.0, 53.0, 54.2, 55.4, 56.0, and 57.0 °C as annealing temperatures, in combination with an initial denaturation at 95 °C for 3 min, followed by 27 cycles of denaturation at 95 °C for 30 s, annealing for 30 s, and extension at 72 °C for 60s. The final PCR assays consisted of 3 min of denaturation at 95 °C, followed by 27 cycles of denaturation at 95 °C for 30 s, annealing at 52 °C for 30 s, and extension at 72 °C for 60 s. The PCR amplicons obtained were visualized through gel electrophoresis using 1.5 % (m/v) agarose gels and performed at 100 V for 1 h. PCR amplicon purification was performed using a Wizard Plus SV Mini-preps DNA purification system (Promega, Madison, Wisconsin, USA). PCR assays for all MCs and food samples were performed in triplicate with sample-specific barcodes (Supplementary Table S2).

### 4.5. PacBio HiFi long-read sequencing

After PCR amplification with the sample-specific-barcoded primers, sequencing library preparation was done following PacBio’s workflow (“Procedure & checklist – Preparing Kinnex libraries from 16S rRNA amplicons”, PacBio Document 103-238-800). Briefly, amplicons were pooled and a Kinnex PCR was performed, by which Kinnex terminal adaptors were added to the amplicons to enable concatenation. Next, amplicons were concatenated using the Kinnex enzyme and ligase. Finally, SMRTbell adapters were ligated to generate circular DNA templates suitable for circular consensus sequencing (CCS). The obtained SMRTbell library was sequenced on a Revio platform (Pacific Biosciences) at the VIB Nucleomics Core Facility (Leuven, Belgium).

### 4.6. Data processing

The reads obtained from the Revio platform were deconcatenated using Skera (v1.3.0, PacBio), and individual amplicons were demultiplexed using Lima (v2.12.0, Pacific Biosciences) with a minimal length cut-off set at 50 base pairs (bp) to avoid removing any short ITS sequences. Demultiplexed reads were further processed using RStudio (version 4.2.1; 58), and amplicon sequence variants (ASVs) were obtained using the DADA2 package (version 1.26.0, 59). The filtering parameters used were minQ = 2, minLen = 300, maxLen = 1000, maxN = 0, and maxEE = 5. Taxonomy was assigned using the UNITE database (V10.0, 60) using a minBoot = 80. Reads not identified at the genus level (‘above genus reads’ or AGR) were filtered out. ASVs were grouped per species, and relative abundances of each species per sample were calculated. Further, the AGR were used as queries for blastn searches using the core_nt database of the National Center for Biotechnology Information (NCBI; July 2025).

Rarefaction curve analysis was performed using the R package vegan (v2.6-4; 61). The divergence rate was calculated as the Bray-Curtis distances between the expected and the observed relative abundance using the same R package. The number of false negatives (FN) was defined as the number of expected taxa that were not recovered, and the false positives (FP) as the number of recovered taxa that were not expected. Furthermore, the number of true positives (TP) was defined as the number of recovered taxa that were expected. Finally, the precision of the method was calculated as TP/(TP+FP) (12).

## Data availability

The sequenced reads are available at the European Nucleotide Archive of the European Bioinformatics Institute (ENA/EBI) under the BioProject accession number PRJEB101075.

## Acknowledgements

The authors would like to thank Stéphane Plaisance, Lim De Swert, and the VIB Nucleomics Core Facility team for the sequencing and technical guidance provided. This work was supported by the Research Council of the Vrije Universiteit Brussel (SRP71). Part of the computational resources used were provided by the Flemish Supercomputer Centre (VSC), funded by the Research Foundation - Flanders (FWO-Vlaanderen) and the Flemish Government. TG is the recipient of a PhD fellowship from the Vrije Universiteit Brussel as part of the European research project HealthFerm, which is co-funded by the European Union under the Horizon Europe grant agreement No. 101060247 and the Swiss State Secretariat for Education, Research and Innovation (SERI) under contract No. 22.00210. Views and opinions expressed are, however, those of the author(s) only and do not necessarily reflect those of the European Union or European Research Executive Agency (REA). Neither the European Union nor REA can be held responsible for them.

## References

(1) Marco ML, Sanders ME, Gänzle M, Arrieta MC, Cotter PD, De Vuyst L, Hill C, Holzapfel W, Lebber S, Merenstein D, Redi G, Wolfe BE, Hutkins R. 2021. The International Scientific Association for Probiotics and Prebiotics (ISAPP) consensus statement on fermented foods. Nat Rev Gastroenterol Hepatol 18:196–208.

(2) Gänzle M. 2022. The periodic table of fermented foods: limitations and opportunities. Appl Microbiol Biotechnol 106: 2815–2826.

(3) Valentino V, Magliulo R, Farsi D, Cotter PD, O’Sullivan O, Ercoloni D, De Filippis F. 2024. Fermented foods, their microbiome and its potential in boosting human health. Microb Biotechnol 17:e14428.

(4) Díaz-Muñoz C, De Vuyst L. 2022. Functional yeast starter cultures for cocoa fermentation. J Appl Microbiol 133:39–66.

(5) Zheng Z, Gänzle M. 2025. Sequence based characterization of microbial communities in food: The panacea for smart detection of food microbes or dirty deeds done dirt cheap? Trends Food Sci Technol 162:105113.

(6) Yap M, Ercolini D, Álvarez-Ordoñez A, O’Toole PW, O’Sullivan O, Cotter PD. 2022. Next-generation food research: use of meta-omic approaches for characterizing microbial communities along the food chain. Annu Rev Food Sci Technol 13:361–384.

(7) Weckx S, Van Kerrebroeck S, De Vuyst L. 2019. Omics approaches to understand sourdough fermentation processes. Int J Food Microbiol 302: 90–102.

(8) Callahan BJ, McMurdie PJ, Rosen MJ, Han AW, Johnson AJA, Holmes SP. 2016. DADA2: high-resolution sample inference from Illumina amplicon data. Nat Methods 13:581–583.

(9) De Filippis F, Laiola M, Blaiotta G, Ercolini D. 2017. Different amplicon targets for sequencing-based studies of fungal diversity. Appl Environ Microbiol 83:e00905–17.

(10) Walsh AM, Crispie F, Claesson MJ, Cotter PD. 2017. Translating omics to food microbiology. Annu Rev Food Sci Technol 8:113–134.

(11) Billington C, Kingsbury JM, Rivas L. 2022. Metagenomics approaches for improving food safety: a review. J Food Prot 85:448–464.

12. Rué O, Coton M, Dugat-Bony E, Howell K, Irlinger F, Legras J-L, Loux V, Michel E, Mounier J, Neuvéglise C, Sicard D. 2023 Comparison of metabarcoding taxonomic markers to describe fungal communities in fermented foods. Peer Community J 3:e97.

(13) Tedersso L, Albertsen M, Anslan S, Callahan B. 2021. Perspectives and benefits of high-throughput long-read sequencing in microbial ecology. Appl Environ Microbiol 87:e00626–21.

(14) Yarza P, Yilmaz P, Pruesse E, Glöckner FO, Ludwig W, Schleifer K-H, Whitman WB, Euzéby J, Amann R, Rosselló-Móra R. 2014. Uniting the classification of cultured and uncultured bacteria and archaea using 16S rRNA gene sequences. Nat Rev Microbiol 12:635–645.

(15) Johnson JS, Spakowicz DJ, Hong B-Y, Petersen LM, Demkowicz P, Chen L, Leopold SR, Hanson BM, Agresta HO, Gerstein M, Sodergren E, Weinstock GM 2019. Evaluation of 16S rRNA gene sequencing for species and strain-level microbiome analysis. Nat Commun 10:5029.

(16) Sun D-L, Jiang X, Wu QL, Zhou N-Y. 2013. Intragenomic heterogeneity of 16S rRNA genes causes overestimation of prokaryotic diversity. Appl Environ Microbiol 79:5962–5969.

(17) Wagner J, Coupland P, Browne HP, Lawley TD, Francis SC, Parkhill J. 2016. Evaluation of PacBio sequencing for full-length bacterial 16S rRNA gene classification. BMC Microbiol 16:274.

(18) Schloss PD, Girard RA, Martin T, Edwards J, Thrash JC. 2016. Status of the archaeal and bacterial census: an update. mBio 7:e00201–16.

(19) Buetas E, Jordán-López M, López-Roldán A, D’Auria G, Martínez-Priego L, De Marco G, Carda-Diéguez M, Mira A. 2024. Full-length 16S rRNA gene sequencing by PacBio improves taxonomic resolution in human microbiome samples. BMC Genomics 25:310.

(20) Callahan BJ, Wong J, Heiner C, Oh S, Theriot CM, Gulati AS, McGill SK, Dougherty MK. 2019. High-throughput amplicon sequencing of the full-length 16S rRNA gene with single-nucleotide resolution. Nucleic Acids Res 47:e103.

(21) Hlangwani E, Abrahams A, Masenya K, Adebo OA. 2023. Analysis of the bacterial and fungal populations in South African sorghum beer (umqombothi) using full-length 16S rRNA amplicon sequencing. World J Microbiol Biotechnol 39:350.

(22) Bongaerts D, Bouchez A, De Roos J, Cnockaert M, Wieme AD, Vandamme P, Weckx S, De Vuyst L. 2024. Refermentation and maturation of lambic beer in bottles: a necessary step for gueuze production. Appl Environ Microbiol 90 :e01869–23.

(23) Jin H, Mo L., Pan L, Hou Q, Li C, Darima I, Yu J. 2018. Using PacBio sequencing to investigate the bacterial microbiota of traditional Buryatian cottage cheese and comparison with Italian and Kazakhstan artisanal cheeses. J Dairy Sci 101:6885–6896.

(24) Decadt H, Weckx S, De Vuyst L. 2023. The rotation of primary starter culture mixtures results in batch-to-batch variations during Gouda cheese production. Front Microbiol 14 :1128394.

(25) Pradal I, González-Alonso V, Wardhana YR, Cnockaert M, Wieme A, Vandamme P, De Vuyst L. 2024. Various cold storage-backslopping cycles show the robustness of *Limosilactobacillus fermentum* IMDO 130101 as starter culture for Type 3 sourdough production. Int J Food Microbiol 411:110522.

(26) Schoch CL, Seifert KA, Huhndorf S, Robert V, Spouge JL, Levesque CA, Chen W, Fungal Barcoding Consortium. 2012. Nuclear ribosomal internal transcribed spacer (ITS) region as a universal DNA barcode marker for Fungi. PNAS 109:6241–6246.

(27) Bokulich NA, Mills DA. 2013. Improved selection of internal transcribed spacer-specific primers enables quantitative, ultra-high-throughput profiling of fungal communities. Appl Environ Microbiol 8:2519–2526.

(28) Hoggard M, Vesty A, Wong G, Montgomery JM, Fourie C, Douglas RG, Biswas K, Taylor MW. 2018. Characterizing the human mycobiota: a comparison of small subunit rRNA, ITS1, ITS2, and large subunit rRNA genomic targets. Front Microbiol 9:2208.

(29) Hu Y, Irinyi L, Hoang MTV, Eenjes T, Graetz A, Stone EA, Meyer W, Schwessinger B, Rathjen JP. 2022. Inferring species compositions of complex fungal communities from long- and short-read sequence data. mBio 13:e02444–21.

(30) Mbarache H, Veillette M, Bilodeau G, Duchaine C. 2020. Comparison of the performance of ITS1 and ITS2 as barcodes in amplicon-based sequencing of bioaerosols. PeerJ 17:e8523.

(31) Tedersoo L, Anslan S, Bahram M, Kõljalg U, Abarenkov K. 2020. Identifying the ‘unidentified’ fungi: a global-scale long-read third-generation sequencing approach. Fungal Divers 103:273–293.

(32) James TY, Marino JA, Perfecto I, Vandermeer J. 2016. Identification of putative coffee rust mycoparasites via single-molecule DNA sequencing of infected pustules. Appl Environ Microbiol 82:e02639–15.

(33) Walder F, Schlaeppi K, Wittwer R, Held AY, Vogelgsang S, van der Heijden MGA. 2017. Community profiling of Fusarium in combination with other plant-associated fungi in different crop species using SMRT sequencing. Front Plant Sci 8:2019.

(34) Latz MAC, Grujcic V, Brugel S, Lycken J, John U, Karlson B, Andersson A, Andersson AF. 2022. Short- and long-read metabarcoding of the eukaryotic rRNA operon: evaluation of primers and comparison to shotgun metagenomics sequencing. Afr J Ecol 22:2304–2318.

(35) Tedersoo L, Tooming-Klunderud A, Anslan S. 2017. PacBio metabarcoding of Fungi and other eukaryotes: errors, biases and perspectives. New Phytol 217:973–976.

(36) Díaz-Muñoz C, Van de Voorde D, Comasio A, Verce M, Hernandez CE, Weckx S, De Vuyst L. 2021. Curing cocoa beans: fine-scale monitoring of the starter culture applied and metabolomics of the fermentation and drying steps. Front Microbiol 11:616875.

(37) González-Alonso V, Pradal I, Wardhana YR, Cnockaert M, Wieme AD, Vandamme P, De Vuyst L. 2024. Microbial ecology and metabolite dynamics of backslopped triticale sourdough productions and the impact of scale. Int J Food Microbiol 408:110445.

(38) Pradal I, González-Alonso V, Wardhana YR, De Vuyst L. 2025. *Companilactobacillus crustorum* LMG 23699 and *Wickerhamomyces anomalus* IMDO 010110 form a candidate, stable, mixed-strain starter culture for sourdough production. Int J Food Microbiol 440:111278.

(39) Čadež N, Poot GA, Raspor P, Smith MT. 2003. *Hanseniaspora meyeri* sp. nov., *Hanseniaspora clermontiae* sp. nov., Hanseniaspora lachancei sp. nov. and Hanseniaspora opuntiae sp. nov., novel apiculate yeast species. Int J Syst Evol Microbiol 53:1671–1680.

(40) Wang Q-M, Li J, Wang S-A, Bai F-Y. 2008. Rapid differentiation of phenotypically similar yeast species by single-strand conformation polymorphism analysis of ribosomal DNA. Appl Environ Microbiol 74:2604–2611.

(41) Heeger F, Bourne EC, Baschien C, Yurkov A, Bink B, Spröer C, Overmann J, Mazzoni CJ, Monaghan MT. 2018. Long-read DNA metabarcoding of ribosomal RNA in analysis of fungi from aquatic environments. Mol. Ecol. Resour. 18:1500–1514.

(42) D’Andreano S, Cuscó A, Francino O. 2020. Rapid and real-time identification of fungi up to species level with long amplicon nanopore sequencing from clinical samples. Biol methods protoc 6:bpaa026.

(43) Krabberød AK, Stokke E, Thoen E, Skrede L, Kauserud H. 2024. The Ribosomal Operon Database: a full-length rDNA operon database derived from genome assemblies. Mol Ecol Resour 25:e14031.

(44) Kurtzman CP. 2003. Phylogenetic circumscription of *Saccharomyces*, *Kluyveromyces* and other members of the Saccharomycetaceae, and the proposal of the new genera *Lachancea*, *Nakaseomyces*, *Naumovia*, *Vanderwaltozyma* and *Zygotorulaspora*. FEMS Yeast Res 4:233–245.

(45) Kurtzman CP, Robnett CJ, Basehoar-Powers E. 2008. Phylogenetic relationships among species of *Pichia*, *Issatchenkia* and *Williopsis* determined from multigene sequence analysis, and the proposal of *Barnettozyma* gen. nov., Lindnera gen. nov. and Wickerhamomyces gen. nov. FEMS Yeast Res 8:939– 954.

(46) Wang L, Groenewald M, Wang QM, Boekhout T. 2015. Reclassification of *Saccharomycodes sinensis*, proposal of *Yueomyces sinensis* gen. nov., comb. nov. within Saccharomycetaceae (Saccharomycetales, Saccharomycotina). PLOS ONE 10:e0136987.

47. Groenewald M, Hittinger CT, Bensch K, Opulente DA, Shen X-X, Li Y, Liu C, LaBella AL, Zhou X, Limtong S, Jindamorakot S, Gonçalves P, Robert V, Wolfe KH, Rosa CA, Boekhout T, Ĉadež N, Péter G, Sampaio JP, Lachance M-A, Yurkov AM, Daniel H-M, Takashima M, Boundy-Mills K, Libkind D, Aoki K, Sugita T, Rokas A. 2023. A genome-informed higher rank classification of the biotechnologically important fungal subphylum Saccharomycotina. Stud Mycol 105:1–22.

(48) Liu F, Hu Z-D, Yurkov A, Chen X-H, Bao W-J, Ma Q, Zhao WN, Pan S, Zhao X-M, Liu J-H, Wang Q-M, Boekhout T. 2024. Saccharomycetaceae: delinaeation of fungal genera based on phylogenomic analyses, genomic relatedness indices and genomics-based synapomorphies. Persoonia 52:1–21.

(49) Bongaerts D, De Roos J, De Vuyst L. 2021. Technological and environmental features determine the uniqueness of the lambic beer microbiota and production process. Appl Environ Microbiol 87:e00612–21.

50. von Gastrow L, Gianotti A, Vernocchi P, Serrazanetti DI, Sicard D. 2023. Chapter 7, Taxonomy, biodiversity, and physiology of sourdough yeasts, p 161–212. In Gobbetti M & Gänzle M (ed), Handbook on Sourdough Biotechnology. Springer Nature, Switzerland.

(51) Urien C, Legrand J, Montalent P, Casaregola S, Sicard D. 2019. Fungal species diversity in French bread sourdoughs made of organic wheat flour. Front Microbiol 10:201.

(52) Comasio A, Harth H, Weckx S, De Vuyst L. 2019. The addition of citrate stimulates the production of acetoin and diacetyl by a citrate-positive *Lactobacillus crustorum* strain during wheat sourdough fermentation. Int J Food Microbiol 289:88–105.

(53) Arora K, Ameur H, Polo A, Di Cagno R, Rizzello CG, Gobbetti M. 2021. Thirty years of knowledge on sourdough fermentation: a systematic review. Trends Food Sci Technol 108:71–83.

(54) Tedersoo L, Lindahl B. 2016. Fungal identification biases in microbiome projects. Environ Microbiol Rep 8:774–779.

(55) Tedersoo L, Anslan S. 2019. Towards PacBio-based pan-eukaryote metabarcoding using full-length ITS sequences. Environ Microbiol Rep 11:659–668.

(56) Bazalová O, Cihlár JZ, Dlouhá Z, Bár L, Dráb V, Kavková M. 2022. Rapid sourdough yeast identification using panfungal PCR combined with high resolution melting analysis. J Microbiol Methods 199:106522.

(57) Sánchez-Adriá I, Sanmartín G, Prieto JA, Esturch F, Fortis E, Randez-Gil F. 2023. Technological and acid stress performance of yeast isolates from industrial sourdough. LWT-Food Sci Technol 184:114957.

58. Posit team. 2025. RStudio: Integrated Development Environment for R. Posit Software, PBC, Boston, MA. http://www.posit.co/.

(59) Callahan BJ, McMurdie PJ, Holmes SP. 2017. Exact sequence variants should replace operational taxonomic units in marker-gene data analysis. ISME J 11:2639–2643.

60. Abarenkov K, Zirk A, Piirmann T, Pöhönen R, Ivanov F, Nilsson RH, Kõljalg U. 2025. UNITE general FASTA release for Fungi. UNITE Community. 10.15156/BIO/3301229

61. (61) Oksanen J, Simpson G, Blanchet F, Kindt R, Legendre P, Minchin P, O’Hara R, Solymos P, Stevens M, Szoecs E, Wagner H, Barbour M, Bedward M, Bolker B, Borcard D, Carvalho G, Chirico M, De Caceres M, Durand S, Evangelista H, FitzJohn R, Friendly M, Furneaux B, Hannigan G, Hill M, Lahti L, McGlinn D, Ouellette M, Ribeiro Cunha E, Smith T, Stier A, Ter Braak C, Weedon J. 2022. vegan: Community Ecology Package. https://CRAN.R-project.org/package=vegan.

